# Transcriptome alterations are enriched for synapse-associated genes in the striatum of subjects with obsessive-compulsive disorder

**DOI:** 10.1101/2020.07.23.216697

**Authors:** Sean C. Piantadosi, Lora L. McClain, Lambertus Klei, Jiebiao Wang, Brittany L. Chamberlain, Sara A Springer, Bernie Devlin, David A. Lewis, Susanne E. Ahmari

## Abstract

**Background:** Obsessive compulsive disorder (OCD) is a chronic and severe psychiatric disorder for which effective treatment options are limited. Structural and functional neuroimaging studies have consistently implicated the orbitofrontal cortex (OFC) and striatum in the pathophysiology of the disorder. Recent genetic evidence points to involvement of components of the excitatory synapse in the etiology of OCD. However, the transcriptional alterations that could link genetic risk to known structural and functional abnormalities remain mostly unknown.

**Methods:** To assess potential transcriptional changes in the OFC and two striatal regions (caudate nucleus and nucleus accumbens) of OCD subjects relative to unaffected comparison subjects, we sequenced messenger RNA transcripts from these brain regions.

**Results:** In a joint analysis of all three regions, 904 transcripts were differentially expressed between 7 OCD versus 8 unaffected comparison subjects. Region-specific analyses highlight a smaller number of differences, which concentrate in caudate and nucleus accumbens. Pathway analyses of the 904 differentially expressed transcripts showed enrichment for genes involved in synaptic signaling, with these synapse-associated genes displaying lower expression in OCD subjects relative to unaffected comparison subjects. Finally, we estimate that cell type fractions of medium spiny neurons are lower whereas vascular cells and astrocyte fractions are higher in tissue of OCD subjects.

**Conclusions:** Together, these data provide the first unbiased examination of differentially expressed transcripts in both OFC and striatum of OCD subjects. These transcripts encode synaptic proteins more often than expected by chance, and thus implicate the synapse as a vulnerable molecular compartment for OCD.

## INTRODUCTION

Obsessive compulsive disorder (OCD), a chronic psychiatric illness, has a lifetime prevalence of 1-3%, affects approximately 50 million people worldwide, and can lead to significant impairment [1–3]. OCD is characterized by intrusive, recurrent thoughts (obsessions) and repetitive behaviors or mental acts (compulsions) that are often performed to reduce the anxiety associated with obsessions [4]. Even with current treatments, many patients continue to experience substantial symptoms and remission is rare [5], underscoring the need for improved understanding of etiology.

Family [6, 7] and twin studies [7] consistently support a genetic contribution to OCD. Heritability estimates of OCD, as well as obsessive-compulsive characteristics, indicate that genetic factors explain between 27%-47% of the phenotypic variance [7–10]. Genetic association studies point to the glutamate transporter gene *SLC1A1* [11–15] and the glutamate receptor genes *GRIN2B* [16, 17] and *GRIK2* [18–20]. Further, genome-wide association studies (GWAS), though underpowered, converge on excitatory synaptic signaling as a potential source of dysfunction [21–23].

Consistent with genetic evidence, neuroimaging research into the neural basis of OCD has highlighted dysfunction within the orbitofrontal cortex (OFC) and striatum, which are connected via dense cortico-striatal glutamatergic projections [4]. Both at rest and following symptom provocation, OCD subjects show hypermetabolism of glucose in the OFC, relative to controls [24–27], and increased blood oxygen-level dependent signals in both OFC and caudate, a subregion of the dorsal striatum [28, 29]. In addition, functional connectivity between OFC and striatum is increased in OCD subjects relative to healthy controls [30, 31] (though see [32]).

Causal links between excitatory synaptic dysfunction at cortico-striatal synapses and OCD-relevant behaviors have also been drawn from rodent studies. Mice with constitutive knockout of *Sapap3* (also known as *Dlgap3*), a post-synaptic density protein enriched at excitatory cortico-striatal synapses, or *Slitrk5*, a post-synaptic transmembrane protein found in excitatory synapses, display compulsive grooming phenotypes and cortical (*Slitrk5-KO*) and striatal (*Sapap3-KO*) hyperactivity that can be rescued with chronic fluoxetine (the first line pharmacotherapy for OCD) treatment [33–35]. Further, these models have altered expression of glutamate receptor transcripts within the striatum, suggesting that removal of critical post-synaptic proteins affects both the structure and function of cortico-striatal synapses and may contribute to compulsive behavior.

Despite these links between genetic risk and neural alterations, and the supporting evidence of causal involvement provided by rodent studies, a dearth of information exists regarding molecular disruptions in these brain regions in OCD. To address this knowledge gap, we recently examined a targeted list of excitatory- and inhibitory-synapse related transcripts in the OFC, caudate, and nucleus accumbens in OCD post-mortem brain tissue. This analysis demonstrated lower excitatory synapse-related transcripts in OCD subjects in both OFC and striatum, though reductions were more pronounced in the OFC [36]. However, this initial study only looked at a small subset of transcripts and therefore could not paint a complete picture of possible OFC and striatal dysfunction. Another recent study, which utilized unbiased RNA-sequencing (RNAseq) to identify gene expression differences in striatal subregions (caudate, putamen, and nucleus accumbens) of OCD subjects compared to unaffected comparison subjects, observed differences in gene expression between striatal subregions and potential involvement of synaptic and immune function protein-encoding transcripts [37]. However, this study only examined the striatum and not its key glutamatergic input, the cortex. Here we used RNA sequencing to examine the transcriptome of two orbitofrontal cortex (medial and lateral OFC) and two striatal subregions (caudate and nucleus accumbens) in OCD subjects and unaffected comparison subjects.

## METHODS

### Human Post-mortem Subjects

Brain specimens (*N*=16) were obtained through the University of Pittsburgh Brain Tissue Donation Program during autopsies conducted by the Allegheny County Medical Examiner’s Office (Pittsburgh, PA) after consent was given by next-of-kin. An independent panel of experienced clinicians reviewed findings from structured interviews with family members, clinical records, toxicology, and neuropathology to make consensus DSM-IV diagnoses. Unaffected comparison subjects underwent identical assessments and were determined to be free of any lifetime psychiatric illnesses. All procedures were approved by the University of Pittsburgh’s Committee for the Oversight of Research and Clinical Training Involving Decedents and Institutional Review Board for Biomedical Research. To reduce biological variance between groups, each subject with OCD (*N*=8) was matched with one unaffected comparison subject for sex (8 males and 8 females), age (standard error of the difference (SED=0.29), post-mortem interval (PMI; SED=3.18), and tissue pH (SED=0.076). The age of unaffected comparison subjects ranged from 20-65 years, while the age of OCD subjects ranged from 20-69 years (Table 1).

**TABLE 1.** Cohort demographics. OCD, obsessive compulsive disorder; PMI, post-mortem interval; RIN, RNA integrity number; SD, standard deviation; n, count; df, degrees of freedom; t, Student’s t-statistic; p, probability.

|  | OCD subjects |  |  | Unaffected comparison subjects |  |  | df | t | p |
| --- | --- | --- | --- | --- | --- | --- | --- | --- | --- |
|  | mean | SD | n | mean | SD | n |  |  |  |
| Age (years) | 47.4 | 14.70 | 7 | 47.6 | 13.42 | 8 | 13 | 0.03 | 0.98 |
| pH | 6.7 | 0.15 | 7 | 6.6 | 0.14 | 8 | 13 | 1.57 | 0.14 |
| Post Mortem Interval<br>(PMI; hours) | 17.8 | 8.37 | 7 | 15.2 | 4.43 | 8 | 13 | 0.75 | 0.46 |
| RNA ratio<br>(260 nm/280 nm) | 1.6 | 0.24 | 7 | 1.5 | 0.31 | 8 | 13 | 0.70 | 0.49 |
| RNA Integrity Number<br>(RIN) | 7.8 | 0.44 | 7 | 7.6 | 0.65 | 8 | 13 | 0.46 | 0.65 |

### Tissue Collection & RNA Extraction

Standardized amounts (50 mm^3^) of gray matter were collected four separate brain regions medial orbitofrontal cortex (mOFC, BA11), lateral orbitofrontal cortex (lOFC, BA47), head of the caudate nucleus, and nucleus accumbens core and RNA was extracted as previously described [36] (Details provided in Supplement 1).

### RNA sequencing

RNA sequencing and quality control (QC) steps are described in detail in Supplement 1. In brief, messenger RNA (mRNA) sequencing was performed at a targeted depth of 40 million reads per sample using the Illumina NextSeq 500 platform (Illumina Inc, San Diego, CA). mRNA reads (4 regions, 16 subjects) were mapped to 58,243 transcripts (Supplemental Figure 1), which were further reduced to 18,993 expressed RefSeq genes (v.2015-01). Of those, genes were retained that had at least 1 count per million in at least half of the samples for at least one brain region; and if, over all subjects and brain regions, gene j’s coefficient of variation CV_j_<T, where T is defined as the mean of CV_j_ over all j plus 3 standard deviations of CV. This resulted in 14,211 genes whose expression passed QC. Next, we evaluated pairwise consensus correlations over genes. Correlation was high between BA11 and BA47, 0.71, compared to others (0.16-0.25; Supplemental Figure 2). Thus, data for BA11 and BA47 were treated as from a single brain region termed “OFC” by averaging expression, per gene and subject, over BA11 and BA47. In this process, we noted one male OCD subject for whom the BA47 and nucleus accumbens samples had likely been inadvertently switched; this subject’s data were removed. After QC, 14,184 genes remained for analyses for all brain regions and 13,623, 13,889, and 13,756 for OFC, caudate, and NAcc, respectively.

### Differential Expression

To identify drivers of variation in gene expression, the relationship between gene expression values and covariates – diagnosis, sex, age, post-mortem interval, pH, RNA integrity number, or brain region – were assessed using generalized linear regression [38] (see Supplement 1 for details).

### Gene Set Enrichment

To investigate whether differentially expressed genes between OCD subjects and unaffected comparison subjects were enriched for biologically relevant gene-set pathways, we used the Gene Set Enrichment Analysis (GSEA) platform [39]. This platform contains various molecular signature databases (MSigDB, v7.0; [40]), including curated gene sets that are publicly available and biologically relevant. Enrichment was evaluated on the differentially expressed genes from the analysis of all brain regions together and per individual brain region (MSigDB sets used are given in Supplement 1).

### Cell Type Composition

The composition of the tissue samples in terms of broad cell type fractions–excitatory neurons, medium spiny neurons (also known as spiny projection neurons), interneurons, astrocytes, oligodendrocytes, ependymal cells, immune cells, and vascular cells – was estimated using deconvolution methods. The principle behind this method is that the bulk expression of a gene for a tissue sample, here measured using RNAseq, is a convolution of the number of cells of each type comprising the sample and the average expression of the gene within the cells of each type. A detailed description of this procedure can be found in [41] and in Supplement 1).

## RESULTS

Here, we highlight two levels of analysis for differentially expressed genes (DEG): from three regions treated jointly (OFC, caudate, and nucleus accumbens), which we refer to as *global analysis*; and from each brain region considered separately, which we refer to as *regional analysis*. We first determined a parsimonious model for gene expression as a function of diagnosis (OCD versus unaffected comparison subjects) and other predictors. For the global analysis of DEG, predictors for the final model included diagnosis and brain region, as well as sex and pH as covariates (Supplemental Table 1). It yielded 904 DEGs (FDR<0.05; 471 were upregulated and 433 were downregulated). Predictors for the regional analyses were the same as the global analysis (e.g. diagnosis, with sex and pH as covariates), although effects were estimated by region. Between diagnostic groups, this analysis identified 29 (23 upregulated and 6 downregulated) and 216 (79 upregulated and 137 downregulated) DEG in the caudate and nucleus accumbens, respectively (FDR<0.05; Figure 1a; Supplemental Table 2), and none for the OFC (Figure S3). Examining overlap of DEG sets between global, caudate, and nucleus accumbens identifies 3 DEG (Figure 1b). There were also 20 DEG in the intersection of the caudate and global gene sets and 177 DEG in the intersection of the nucleus accumbens and global gene sets (Figure 1b). Interestingly, no overlapping DEG were detected between the caudate and the nucleus accumbens.

**FIGURE 1.**
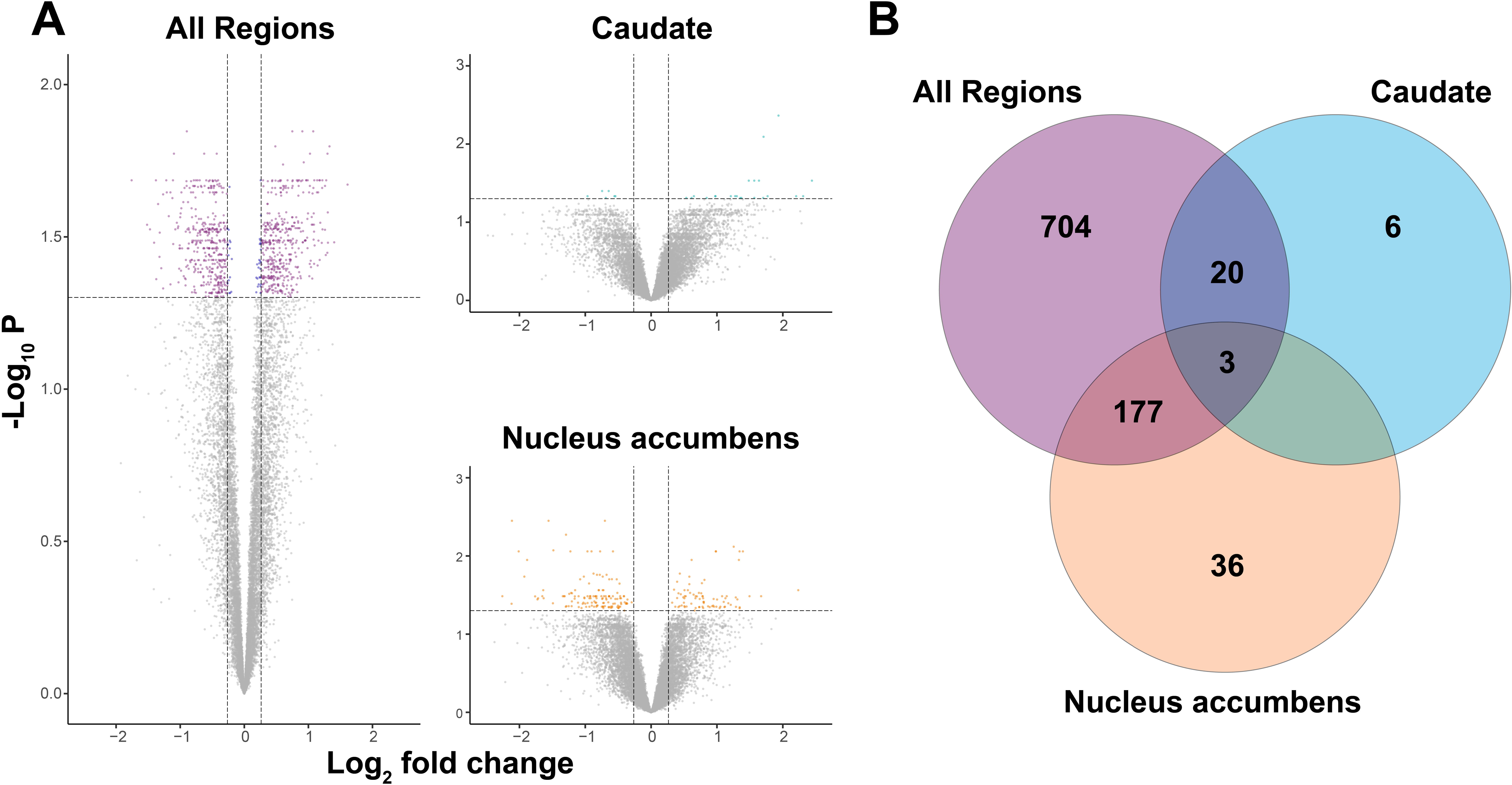
Volcano plots of differentially expressed genes between obsessive compulsive disorder (OCD) subjects and unaffected comparison subjects. RNA sequencing was performed on post-mortem brain tissue originating from Brodmann areas 11 and 47 (OFC), the caudate, and the nucleus accumbens of 7 OCD subjects and 8 unaffected comparison subjects. **(A)** Left panel: Differentially expressed genes were determined for all brain regions analyzed together. Upper right panel: Differentially expressed genes in caudate. Lower right panel: Differentially expressed genes in nucleus accumbens. The y-axis represents the (−log_10_P-value) and the x-axis represents the gene expression log_2_fold change. Vertical dashed lines (±0.26 log_2_ fold change) indicate gene expression differences between OCD subject and unaffected comparison subject cohorts, where upregulated genes are positive and downregulated genes are negative. The horizontal line demarcates significantly different gene expression differences between OCD subjects and unaffected comparison subjects (false discovery rate q-value <0.05; purple (≥0.26 or ≤−0.26 log_2_ fold change) / blue (≤0.26 and ≥−0.26 log_2_ fold change) dots = significant, gray dots = non-significant). Note: the volcano plot for the cortical regions (Brodmann areas 11 and 47) is shown in Figure S3. **(B)** Venn diagram depicting the overlapping differentially expressed genes between all brain regions (purple), caudate (cyan), and nucleus accumbens (orange). There were no overlapping differentially expressed genes at the intersection between caudate and nucleus accumbens.

### Gene Set Enrichment

DEG sets were next assessed for enrichment of gene functions (FDR adjusted for multiple testing, both here and for enrichment analysis). To do so, the 904 (global), 29 (caudate), and 216 (nucleus accumbens) DEG sets were queried against the MSigDB C2 curation, the C5 curation (GO sets), and the Hallmark gene sets. The global DEG were significantly enriched for 13 Reactome, 32 GO, and 5 Hallmark gene sets. DEG from the nucleus accumbens were enriched for one Hallmark gene set, and DEG from the caudate were enriched for one REACTOME and two GO gene sets after correction for multiple testing (Tables S3a/S3b).

For the global analysis, we noted that several of the enriched GO gene set pathways had a high proportion of overlapping genes. We leveraged this overlap to explore the links between enriched gene sets and draw broader conclusions about the nature of transcription alterations in OCD. Hierarchical clustering revealed four distinct sub-networks of gene sets. Of note, cluster 1 contained 16 gene sets that could be attributed to the synapse (Figure 2). We also tabulated the number of enriched gene sets with either upregulated or downregulated DEG in OCD subjects versus unaffected comparison subjects (Table S3). Because our previous qPCR work [36] identified lower levels of several synaptic transcripts within the OFC and striatum of OCD subjects compared to unaffected comparison subjects, we sought to determine whether the genes identified in these synaptic gene sets contained more downregulated genes than other clusters. For the 16 synaptic gene sets identified in cluster 1, the fraction of genes that were downregulated for each set was significantly higher than that for the remaining 16 gene sets (mean fraction of downregulated genes was 0.712 and 0.507, respectively; t=6.26; df=30; p=6.7×10^−7^). Furthermore, we found that there were significant differences in the mean fraction of downregulated genes across the four clusters identified in Figure 2 (analysis of variance, F[3, 28]=28.46, p=1.19×10^−8^). The mean fraction of downregulated genes for clusters 1, 2, 3, and 4 was 0.76, 0.63, 0.69, and 0.38, respectively (Figure 2; Table S3): cluster 1, which is comprised of mostly synaptic neurotransmission enriched gene sets, had a significantly higher fraction of down-regulated genes (t=4.91, Bonferroni corrected p=1.34×10^−4^), while cluster 4, containing pathways involved in receptor tyrosine kinase activity, circulatory system development, and plasma membrane regulation, had a significantly higher fraction of upregulated genes (t=−7.4, Bonferroni corrected p=4.94×10^−7^). Clusters 2 and 3, representing cytosolic plasma membrane components and transmembrane transport, respectively, had more similar levels of up and downregulated genes.

**FIGURE 2.**
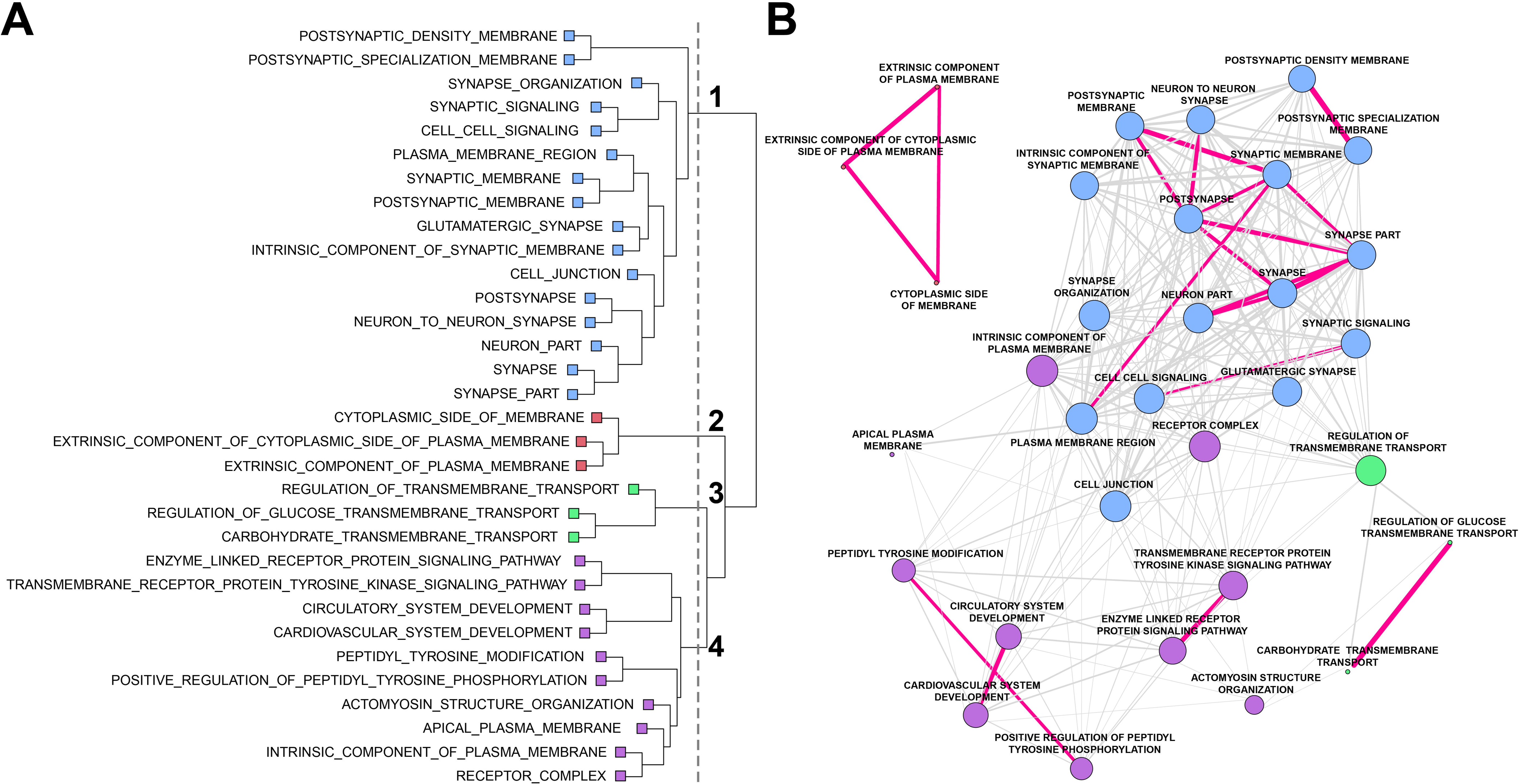
Gene set enrichment analysis for obsessive compulsive disorder (OCD). Following RNA sequencing, global differentially expressed genes were determined between OCD subjects and unaffected comparison subjects (904 genes). Significant gene sets from the gene ontology (GO) pathways were determined using Fisher’s exact test. **(a)** Gene co-occurrence cluster dendrogram for gene set enrichment in OCD. Following gene set enrichment analysis, the number of co-occurring genes among the GO pathways were assessed. Four main branches were identified on the basis of the cut-point (vertical gray dashed line) established by visual inspection and color-coded per branch (blue: cluster 1; red: cluster 2, green: cluster 3, purple: cluster 4). **(b)** Cluster plot of gene sets associated with OCD. Genes were assessed using hierarchical clustering on the distances computed per gene set. The edges (lines) connecting nodes indicate the number of co-occurring OCD genes, where magenta lines have at least half of the genes co-occurring between the nodes (gene pathways), and gray lines indicate less than half of the OCD genes were co-occurring. Co-occurring genes were defined as the fraction of genes present in two pathways given the total number of unique genes represented in both pathways. The thickness of the magenta lines indicates more (thicker) or less (thinner) gene co-occurrence between two given gene sets. Note, the length of connecting vertices does not indicate distance between nodes.

### Cell-type representation within tissue

We used the gene expression patterns in the bulk tissue for each of the three brain regions to estimate the fractions of eight broad cell types: excitatory neurons, inhibitory interneurons, medium spiny neurons (a.k.a. spiny projection neurons), astrocytes, ependymal cells, immune cells, oligodendrocytes, and vascular cells (Table S4). After adjusting for covariates, OCD subjects were significantly different from unaffected comparison subjects for the representation of the following cell types in the global analysis: astrocytes (p=0.005), vascular cells (p=0.002), and medium spiny neurons (p=0.009; Figure 3a).The remaining cell types were not significantly different. Astrocyte cell fractions were larger in OCD subject samples, as were vascular cell fractions, whereas medium spiny neuron fractions were smaller in OCD samples (Figure 3a). Regional analyses suggest smaller interneuron fractions in the OFC (Figure 3b), smaller medium spiny neuron fractions in the caudate (Figure. 3c), smaller ependymal cell fractions in the nucleus accumbens (Figure 3d), and a larger fraction of vascular cells in the OFC of OCD subjects compared to unaffected comparison subjects (Figure 3b; Table S5). While regional results do not achieve significance after correction for multiple testing, they are consistent with global results.

**FIGURE 3.**
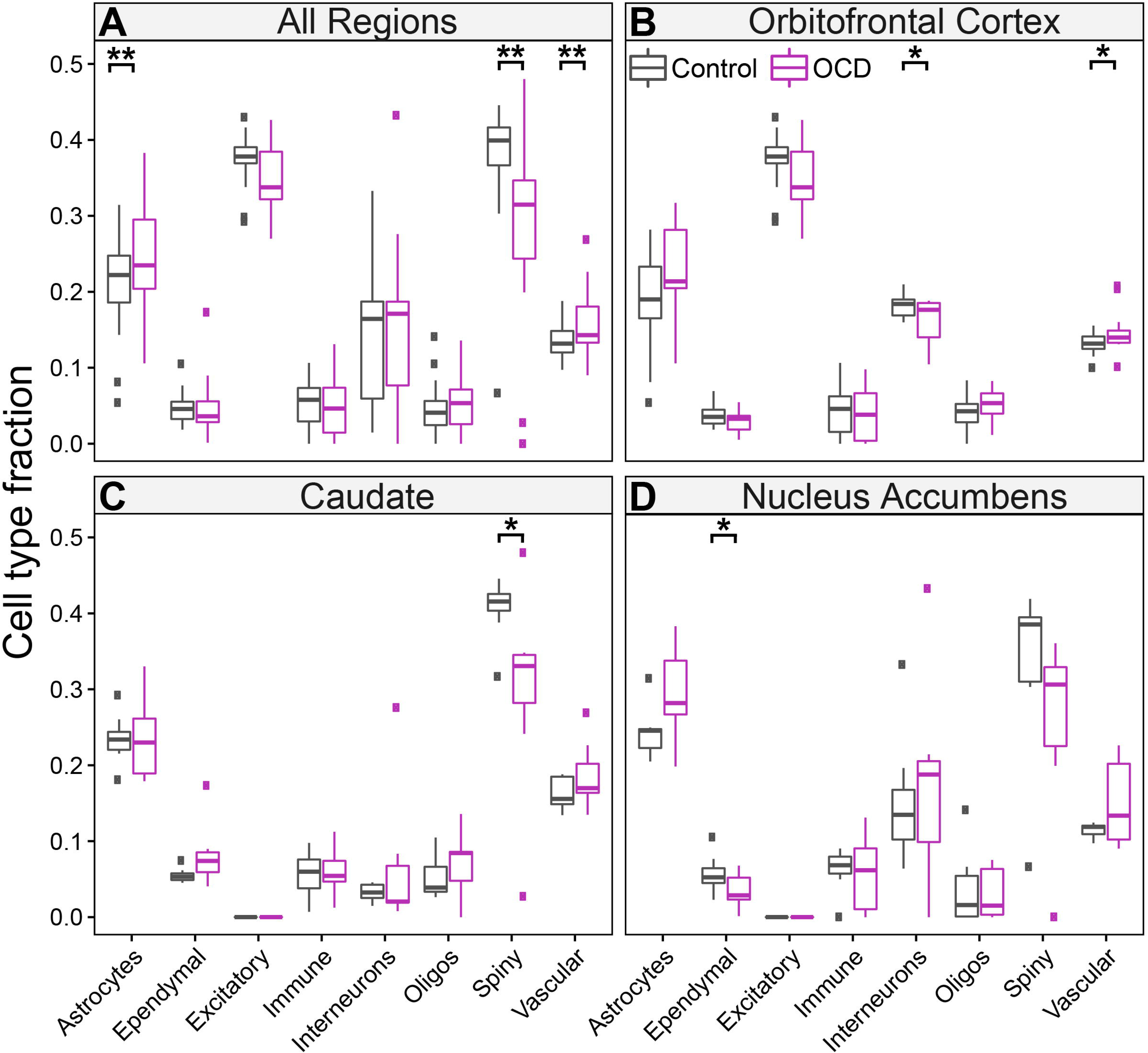
Cell type fractions in OCD subjects and unaffected comparison subjects. Cell type fractions were determined across all brain regions for OCD subjects and unaffected comparison subjects separately. Gene expression patterns in single-cell data were used to estimate the fraction of eight broadly defined cell types: astrocytes, ependymal cells, excitatory neurons, immune cells, interneurons, oligodendrocytes, medium spiny neurons (a.k.a. spiny projection neurons), and vascular cells. Linear regression was used to delineate OCD subjects (purple) from unaffected comparison subjects (grey) in each of the eight cell fractions. Boxplots are displayed, where ** indicates p<0.01 and * indicates p<0.05 (uncorrected for multiple comparisons). The mean cell type fractions were computed based on all four brain regions; however, when the cell type did not exist in the brain region, it was not included in the calculation. Thus, the mean cell type fraction for the excitatory neurons was based only on the cortical regions and the mean cell type fraction for spiny projection neurons was based on the caudate and nucleus accumbens regions.

## DISCUSSION

Although the etiology of OCD remains unknown, convergent evidence suggests disruptions in critical components of cortico-striatal glutamatergic synapses [7, 21, 22, 36, 37, 42]. However, no comprehensive analysis of the OCD transcriptome has been performed simultaneously in tissue from OFC and striatum, the most commonly identified regions with functional and structural alterations in neuroimaging studies [31, 43–47]. We therefore examined the transcriptome in two orbitofrontal (medial, lateral) and two striatal (caudate, nucleus accumbens) brain regions in subjects with OCD and matched unaffected comparison subjects. We first looked at global differential expression in all regions jointly and identified 904 transcripts that were differentially expressed as a function of OCD diagnosis (Figure 1a, *left*). Interestingly, most of DEG were found in the caudate and nucleus accumbens (Figure 1a, *right*) and not in OFC (Figure S3). Gene set enrichment and hierarchical clustering of all differentially expressed transcripts identified a hub of pathways involved in synaptic neurotransmission and associated with the glutamatergic synapse (Figure 2). Together, these data further support the involvement of cortical and striatal glutamatergic synaptic dysfunction in OCD pathogenesis. Using our global DEG data, we also estimated broad cell type representation from our tissue samples, finding significantly different fractions of astrocytes, vascular cells, and medium spiny neurons between OCD and unaffected comparison subjects (Figure 3).

Correlational evidence of structural and functional changes in both the OFC and the striatum in OCD has accumulated over the past 30 years [4]. More recent rodent studies have functionally linked these regions to OCD-relevant behavior [48] and highlighted a critical role for excitatory synaptic signaling through the use of transgenic knockout mice lacking proteins involved in normal excitatory synaptic neurotransmission [34, 35, 49–51]. Furthermore, genetic studies of OCD subjects frequently identify associations with genes involved in excitatory synaptic signaling [11, 12, 16, 17, 19–22, 52]. Consistent with these findings, we observed alterations of transcripts involved in synaptic signaling in OCD subjects via gene-set analysis. Of the 32 significant gene-sets we identified, 16 were classified as involved in synaptic neurotransmission (Figure 2). Genes in these sets were disproportionately downregulated (Table S3b), consistent with our previous qPCR findings in the tissue from these same subjects [36] and with genetic knockout mice that display compulsive behavior [35, 49]. Future molecular studies must investigate how downregulation of synaptic gene expression could lead to observed hyperactivity in the OFC and striatum of OCD patients. In the OFC, hyperactivity may be driven through a decrease in expression patterns representative of inhibitory interneurons (Figure 3b), which regulate excitatory/inhibitory balance. In the striatum, downregulation of synaptic gene expression is likely related to a decrease in the expression profile of medium spiny neurons (Figure 3a,c,d), which are the primary recipients of cortico-striatal glutamatergic input. It will be of interest to determine whether alterations in cell type expression profiles are due to a change in the absolute number of that cell type in OCD subjects, a shift in an individual cell’s expression profile (e.g. a decrease in the number of synapses), or compensatory changes due to alterations in other cell types.

In addition to strengthening the link between OCD pathophysiology and synaptic dysfunction in cortical and striatal brain regions, RNAseq and enrichment analysis allowed us to identify novel signaling pathways that may be disrupted in OCD subjects (Figure 2). We find that gene sets related to cytosolic membrane components (cluster 2) and transmembrane transport (cluster 3) contain similar numbers of up and downregulated gene sets, suggesting complex differential regulation between OCD and unaffected comparison subjects. Cluster 2 pathways are comprised of genes encoding cytosolic proteins associated with receptors, such as G proteins and regulatory subunits, as well as other cytosolic components that are critical for intracellular signaling and interact with proteins found in cluster 1. Cluster 3 contains genes that regulate glucose and carbohydrate transport and may indicate different metabolic needs in the OFC and striatum of OCD subjects. This would be consistent with human imaging studies finding hyperactivity and increased glucose metabolism in these regions in OCD [24–26], and could be related to the changes in cell type expression profiles observed in our cell type deconvolution. For example, the metabolic needs of astrocytes, whose expression profile is increased in OCD (Figure 3a), differ from that of neurons [53].

Compared to synaptic enrichment (cluster 1), which included mostly downregulated DEG, cluster 4 pathways associated with circulatory and cardiovascular system development, transmembrane tyrosine kinase activity (RTK), and components of the plasma membrane, mostly consisted of upregulated DEG (Table S3). These upregulated gene sets are consistent with the increased vascular cell expression profile observed in our cell type deconvolution (Figure 3a), and increased angiogenesis in these regions may play a role in the elevated blood oxygen level dependent (BOLD) signal observed in functional imaging studies of OCD subjects. While previous studies of RTK alterations in OCD have identified an association between tropomyosin receptor kinase B/neurotrophic tyrosine kinase receptor type 2 (*TRKB/NTRK2*) and OCD [52], our data suggest a role for other kinases, including Janus kinase 3 (*JAK3*) and protein kinase C beta (*PRKCB1*), though their precise function remains to be determined.

To date, there remains a dearth of studies using an unbiased approach to examine the molecular underpinnings of OCD and related disorders, likely due to the difficulty of obtaining high quality post-mortem tissue samples. In the first transcriptomic study of OCD tissue, Jaffe and colleagues [54] used a microarray approach to examine DEG in the dlPFC as a function of several disorders typified by obsessive thoughts and compulsive behaviors (eating disorders, OCD, OCPD, and tic disorder). The authors found 286 DEG when comparing OCD/OCPD/tic cases to controls. Interestingly, out of the 904 global DEG in our study, only 13 were identified as differentially expressed by Jaffe et al [54]. While this is consistent with statistical challenges owing to the relatively small sample size for both studies (see Fromer, Roussos [55]), it could also indicate that molecular dysregulation in the dlPFC could be distinct from what is seen in the adjacent OFC, as well as the striatum. This is especially interesting in light of neuroimaging findings that suggest hyperactivity at rest in the OFC [24, 25, 56, 57], while no such evidence exists for the dlPFC (though lateral OFC and dlPFC hypoactivity have been consistently observed in OCD patients during cognitive tasks [58, 59]). Studies directly comparing differential expression in the dlPFC and OFC of the same subjects may clarify whether gene expression changes are global or region specific.

A second recent study by Lisboa and colleagues [37] utilized RNAseq to assess gene expression in three striatal regions (caudate, putamen, and nucleus accumbens) in OCD and comparison subjects [37]. Our findings of differences in gene expression within striatal subregions, with no genes identified as differentially expressed in both the caudate and the nucleus accumbens (Figure 1b), are broadly consistent with this work. The authors also identified gene set enrichment of synaptic related transcripts, supporting our observation of functional enrichment of highly correlated synaptic gene sets (Figure 2; Table S3a) that could be primarily driven by changes in striatal gene expression. A key difference between our findings and those of Lisboa et al. is that our analyses did not reveal any changes in immune- or microglia-related gene sets. There are many possible explanations for this, including the age of subjects (age range 58-98 in Lisboa et al. [37], age range 20-69 in the current work), as aging is associated with changes in immune system gene expression [60]. The role of the immune system in OCD (and its interaction with age and disease length) requires further study.

Because post-mortem studies of psychiatric disease have typically relied on whole tissue homogenates [61], they have traditionally been agnostic about the potential contributions of genetically and phenotypically distinct cell types to disease pathology. Here we addressed this limitation by utilizing a deconvolutional approach to estimate cell type specific fractions of whole tissue homogenates from OCD subjects and unaffected comparison subjects. Importantly, this analysis does not differentiate whether differences in cell type specific fractions are due to changes in absolute cell number or changes in the transcriptional activity of single cells. We do observe that OCD subjects have lower cell type expression profiles of medium spiny neurons, the predominant cell type in the striatum [62], pointing us to future studies to discriminate between these two possibilities. Interestingly, we also observed increases in astrocyte and vascular cell type fractions in OCD subjects. Though there is limited evidence linking these cell types to OCD, it was recently demonstrated that altering astrocytic calcium-dependent signaling in mice produces an OCD-like compulsive grooming phenotype and striatal microcircuit deficits [63]. Further, astrocytes and vascular cells interact at tight junctions to regulate blood-brain barrier function. An increase in these cell fractions might reflect a compensatory change due to the hypermetabolism commonly observed in both the OFC and striatum of OCD subjects [25, 64–66].

It is important to note several limitations of the present study and the measures we have taken to account for them. First, our cohort is small, containing only seven OCD subjects and eight unaffected comparison subjects. Obtaining post-mortem tissue with sufficiently high RNA quality to perform robust RNA-sequencing (Table 1; Figure S1) remains a challenge and because of this it is difficult to evaluate the effect of all potential covariates on gene expression. Here we built a model including only covariates that statistically affected transcript expression (see Methods). A second related point is that DEG may be detected due to several factors, including (1) large case versus unaffected subject differences over all three regions, (2) large differences within only one region, (3) or modest but consistent differences for all three regions, among other scenarios. Global analysis could detect DEG for any of these scenarios, within the limits of statistical power, whereas regional analysis would likely detect DEG only for (1) or (2). If (3) would be a better model for most DE genes, global analysis has better power. Third, although these data are broadly consistent with our previous report of down-regulation of a small targeted set of glutamatergic transcripts assessed via qPCR across OFC and striatal brain regions in these same OCD subjects [36], here we did not observe significant DEG in the OFC alone. Notably, the normalized qPCR expression from [36] and the RNAseq gene expression levels reported here were significantly positively correlated as expected (r=0.302, p<0.0001, data not shown). It is therefore likely that there are changes in transcript levels in the OFC of our OCD subjects relative to unaffected comparison subjects, but our small sample size and multiple comparison correction limited our ability to detect them. This interpretation is bolstered by our global analysis demonstrating many more significant differentially expressed transcripts in the global condition (OFC and striatum) than in the striatum alone. Despite these limitations, the utility of performing unbiased transcriptomic analysis is clear as it provides a more complete picture of dysfunction across regions and allows for the identification of novel transcripts and signaling pathways that would not otherwise be detected.

Here we conducted the first unbiased transcriptomic analysis of the OFC and striatum of post-mortem tissue from OCD and unaffected comparison subjects. We found significantly lower expression of transcripts associated with synaptic neurotransmission across all regions and determined that these changes could be strongest in medium spiny neurons in the caudate nucleus and nucleus accumbens. Further examination of these data could help identify novel treatment targets for OCD and related disorders and will serve as a foundation for future larger studies.

## ACKNOWLEDGEMENTS

The authors would also like to thank the patients and their families for their generous tissue donation. Tissue from some subjects was obtained from the NIH NeuroBioBank at the University of Pittsburgh Brain Tissue Donation Program. We would also like to thank Kelly Rogers and Dr. Jill Glausier for their assistance with subject selection and tissue blocking and cutting, as well as Dr. Sue Johnston for her assistance with clinical assessments of post-mortem subjects.

This work was supported by National Institute of Mental Health (NIMH) Grant MH104255-04 (SEA), the Brain Research Foundation Fay and Frank Seed Grant (SEA, SCP), the International Obsessive Compulsive Disorder Foundation (IOCDF) Breakthrough Award (SEA), National Institute of Health (NIH) Institutional Training Grant T32NS007433-19 (SCP, SAS), National Institute of Drug Abuse (NIDA) Training Grant T32DA031111 (SCP), National Institute of General Medical Sciences of the NIH under Award Number T32GM008208 (BLC), and an Achievement Rewards for College Scientists (ARCS) Pittsburgh Chapter Award (SAS).

---

**SUPPLEMENTAL TABLE 1.** <u>Number of genes for which the Bayesian information criterion (BIC) improved during model selection</u>.

**SUPPLEMENTAL TABLE 2.** <u>Analysis of differential gene expression for combined brain regions and for each brain region separately</u>. FC, fold change; CPM, counts per million; t, Student’s t-statistic; FDR, false discovery rate.

**SUPPLEMENTAL TABLE 3.** <u>Gene sets enriched for differentially expressed genes between OCD subjects and unaffected comparison subjects</u>. Gene sets enriched for differentially expressed genes were identified using Fisher’s exact test. Gene sets containing at least 30 genes were considered significant if the enrichment odds ratio was >1.0 and the False Discovery Rate (FDR) q-value was <0.05. A summary of the enriched gene sets is provided in Supplemental Table 3a. Enriched gene sets were identified that contained either upregulated or downregulated genes in OCD subjects versus unaffected comparison subjects (Supplemental Table 3b); that contained only upregulated genes (Supplemental Table 3c); or that contained only downregulated genes (Supplemental Table 3d). DEG: differentially expressed genes; OR: odds ratio; P: probability.

**SUPPLEMENTAL TABLE 4.** <u>Cell-type fraction estimations in post-mortem brain tissue from OCD subjects and unaffected comparison subjects</u>.

**SUPPLEMENTAL TABLE 5** <u>Region-specific differences of cell type fractions between OCD subjects and unaffected comparison subjects</u>. Listed numbers = p-values (uncorrected for multiple comparisons). As expected, there were zero estimated fractions for medium spiny neurons in the orbitofrontal cortex and for excitatory neurons in caudate and nucleus accumbens; thus, the corresponding p-values are marked NA.

**SUPPLEMENTAL FIGURE 1.** <u>RNA sequencing data read counts</u>. The counts of the number of RNA sequencing reads that mapped (blue) or did not map (red) to the human reference sequence are shown per subject per brain region. X-axis indicates subject number.

**SUPPLEMENTAL FIGURE 2.** <u>Hierarchical clustering and heatmap of RNA expression obtained from four post-mortem brain regions in seven OCD subjects and eight unaffected comparison subjects</u>. The similarity between each sample pair was computed using the Euclidean distance of the log_2_(count per million) reads. Two main clusters were observed containing the Brodmann areas (pink and gray boxes indicated at the top of the figure) and the striatal regions (blue and orange boxes indicated at the top of the figure).

**SUPPLEMENTAL FIGURE 3.** <u>Volcano plot of the differentially expressed genes from cortical regions in OCD subjects</u>. The y-axis represents the (−log_10_P-value) and the x-axis represents the gene expression log_2_fold change. Vertical dashed lines (±0.26 log_2_ fold change) indicate expression difference where upregulated genes are positive and downregulated genes are negative. The horizontal line demarcates significantly different gene expression differences between OCD subjects and unaffected comparison subjects (false discovery rate q-value <0.05).

## LITERATURE CITED

1. Koran, L.M., et al., Practice guideline for the treatment of patients with obsessive-compulsive disorder. Am J Psychiatry, 2007. 164(7 Suppl): p. 5–53.

2. Koran, L.M., M.L. Thienemann, and R. Davenport, Quality of life for patients with obsessive-compulsive disorder. Am J Psychiatry, 1996. 153(6): p. 783–8.

3. Koran, L.M., Quality of life in obsessive-compulsive disorder. Psychiatr Clin North Am, 2000. 23(3): p. 509–17.

4. Pauls, D.L., et al., Obsessive-compulsive disorder: an integrative genetic and neurobiological perspective. Nat Rev Neurosci, 2014. 15(6): p. 410–24.

5. Pittenger, C., et al., Clinical treatment of obsessive compulsive disorder. Psychiatry (Edgmont), 2005. 2(11): p. 34–43.

6. Geller, D.A., Obsessive-compulsive and spectrum disorders in children and adolescents. Psychiatr Clin North Am, 2006. 29(2): p. 353–70.

7. Pauls, D.L., The genetics of obsessive-compulsive disorder: a review. Dialogues Clin Neurosci, 2010. 12(2): p. 149–63.

8. Bolton, D., et al., Obsessive-compulsive disorder, tics and anxiety in 6-year-old twins. Psychol Med, 2007. 37(1): p. 39–48.

9. Tambs, K., et al., Structure of genetic and environmental risk factors for dimensional representations of DSM-IV anxiety disorders. Br J Psychiatry, 2009. 195(4): p. 301–7.

10. van Grootheest, D.S., et al., Twin studies on obsessive-compulsive disorder: a review. Twin Res Hum Genet, 2005. 8(5): p. 450–8.

11. Arnold, P.D., et al., Glutamate transporter gene SLC1A1 associated with obsessive-compulsive disorder. Arch Gen Psychiatry, 2006. 63(7): p. 769–76.

12. Dickel, D.E., et al., Association testing of the positional and functional candidate gene SLC1A1/EAAC1 in early-onset obsessive-compulsive disorder. Arch Gen Psychiatry, 2006. 63(7): p. 778–85.

13. Samuels, J., et al., Comprehensive family-based association study of the glutamate transporter gene SLC1A1 in obsessive-compulsive disorder. Am J Med Genet B Neuropsychiatr Genet, 2011. 156B(4): p. 472–7.

14. Shugart, Y.Y., et al., A family-based association study of the glutamate transporter gene SLC1A1 in obsessive-compulsive disorder in 378 families. Am J Med Genet B Neuropsychiatr Genet, 2009. 150B(6): p. 886–92.

15. Stewart, S.E., et al., Association of the SLC1A1 glutamate transporter gene and obsessive-compulsive disorder. Am J Med Genet B Neuropsychiatr Genet, 2007. 144B(8): p. 1027–33.

16. Arnold, P.D., et al., Association of a glutamate (NMDA) subunit receptor gene (GRIN2B) with obsessive-compulsive disorder: a preliminary study. Psychopharmacology (Berl), 2004. 174(4): p. 530–8.

17. Kohlrausch, F.B., et al., Association of GRIN2B gene polymorphism and Obsessive Compulsive disorder and symptom dimensions: A pilot study. Psychiatry Res, 2016. 243: p. 152–5.

18. Delorme, R., et al., Frequency and transmission of glutamate receptors GRIK2 and GRIK3 polymorphisms in patients with obsessive compulsive disorder. Neuroreport, 2004. 15(4): p. 699–702.

19. Rajendram, R., et al., Glutamate Genetics in Obsessive-Compulsive Disorder: A Review. J Can Acad Child Adolesc Psychiatry, 2017. 26(3): p. 205–213.

20. Sampaio, A.S., et al., Association between polymorphisms in GRIK2 gene and obsessive-compulsive disorder: a family-based study. CNS Neurosci Ther, 2011. 17(3): p. 141–7.

21. Stewart, S.E., et al., Genome-wide association study of obsessive-compulsive disorder. Mol Psychiatry, 2013. 18(7): p. 788–98.

22. Mattheisen, M., et al., Genome-wide association study in obsessive-compulsive disorder: results from the OCGAS. Mol Psychiatry, 2015. 20(3): p. 337–44.

23. (OCGAS), I.O.C.D.F.G.C.I.-G.a.O.C.G.A.S., Revealing the complex genetic architecture of obsessive-compulsive disorder using meta-analysis. Mol Psychiatry, 2018. 23(5): p. 1181–1188.

24. Baxter, L.R., et al., Local cerebral glucose metabolic rates in obsessive-compulsive disorder. A comparison with rates in unipolar depression and in normal controls. Archives of general psychiatry, 1987. 44(3): p. 211–218.

25. Baxter, L.R., Jr., et al., Cerebral glucose metabolic rates in nondepressed patients with obsessive-compulsive disorder. Am J Psychiatry, 1988. 145(12): p. 1560–3.

26. Swedo, S.E., et al., Cerebral glucose metabolism in childhood-onset obsessive-compulsive disorder. Archives of general psychiatry, 1989. 46(6): p. 518–523.

27. Rauch, S.L., et al., Regional cerebral blood flow measured during symptom provocation in obsessive-compulsive disorder using oxygen 15-labeled carbon dioxide and positron emission tomography. Arch Gen Psychiatry, 1994. 51(1): p. 62–70.

28. Breiter, H.C., et al., Functional magnetic resonance imaging of symptom provocation in obsessive-compulsive disorder. Arch Gen Psychiatry, 1996. 53(7): p. 595–606.

29. Adler, C.M., et al., fMRI of neuronal activation with symptom provocation in unmedicated patients with obsessive compulsive disorder. J Psychiatr Res, 2000. 34(4-5): p. 317–24.

30. Abe, Y., et al., Hyper-influence of the orbitofrontal cortex over the ventral striatum in obsessive-compulsive disorder. Eur Neuropsychopharmacol, 2015. 25(11): p. 1898–905.

31. Harrison, B.J., et al., Altered corticostriatal functional connectivity in obsessive-compulsive disorder. Arch Gen Psychiatry, 2009. 66(11): p. 1189–200.

32. Posner, J., et al., Reduced functional connectivity within the limbic cortico-striato-thalamo-cortical loop in unmedicated adults with obsessive-compulsive disorder. Hum Brain Mapp, 2014. 35(6): p. 2852–60.

33. Welch, J.M., et al., Cortico-striatal synaptic defects and OCD-like behaviours in Sapap3-mutant mice. Nature, 2007. 448(7156): p. 894–900.

34. Burguiere, E., et al., Optogenetic stimulation of lateral orbitofronto-striatal pathway suppresses compulsive behaviors. Science, 2013. 340(6137): p. 1243–6.

35. Shmelkov, S.V., et al., Slitrk5 deficiency impairs corticostriatal circuitry and leads to obsessive-compulsive-like behaviors in mice. Nat Med, 2010. 16(5): p. 598–602, 1p following 602.

36. Piantadosi, S.C., et al., Lower excitatory synaptic gene expression in orbitofrontal cortex and striatum in an initial study of subjects with obsessive compulsive disorder. Mol Psychiatry, 2019.

37. Lisboa, B.C.G., et al., Initial findings of striatum tripartite model in OCD brain samples based on transcriptome analysis. Sci Rep, 2019. 9(1): p. 3086.

38. Bates, D., et al., Fitting Linear Mixed-Effects Models Using lme4. 2015, 2015. 67(1): p. 48 %J Journal of Statistical Software.

39. Subramanian, A., et al., Gene set enrichment analysis: a knowledge-based approach for interpreting genome-wide expression profiles. Proc Natl Acad Sci U S A, 2005. 102(43): p. 15545–50.

40. Liberzon, A., et al., Molecular signatures database (MSigDB) 3.0. Bioinformatics, 2011. 27(12): p. 1739–40.

41. Wang, J., B. Devlin, and K. Roeder, Using multiple measurements of tissue to estimate subject-and cell-type-specific gene expression. Bioinformatics, 2020. 36(3): p. 782–788.

42. Pittenger, C., M.H. Bloch, and K. Williams, Glutamate abnormalities in obsessive compulsive disorder: neurobiology, pathophysiology, and treatment. Pharmacol Ther, 2011. 132(3): p. 314–32.

43. Menzies, L., et al., Neurocognitive endophenotypes of obsessive-compulsive disorder. Brain, 2007. 130(Pt 12): p. 3223–36.

44. Menzies, L., et al., Integrating evidence from neuroimaging and neuropsychological studies of obsessive-compulsive disorder: the orbitofronto-striatal model revisited. Neuroscience and biobehavioral reviews, 2008. 32(3): p. 525–549.

45. Rauch, S.L., et al., Probing striatal function in obsessive-compulsive disorder: a PET study of implicit sequence learning. J Neuropsychiatry Clin Neurosci, 1997. 9(4): p. 568–73.

46. Rotge, J.Y., et al., Meta-analysis of brain volume changes in obsessive-compulsive disorder. Biol Psychiatry, 2009. 65(1): p. 75–83.

47. Saxena, S. and S.L. Rauch, Functional neuroimaging and the neuroanatomy of obsessive-compulsive disorder. Psychiatr Clin North Am, 2000. 23(3): p. 563–86.

48. Ahmari, S.E., et al., Repeated cortico-striatal stimulation generates persistent OCD-like behavior. Science, 2013. 340(6137): p. 1234–9.

49. Welch, J.M., et al., Cortico-striatal synaptic defects and OCD-like behaviours in Sapap3-mutant mice. Nature, 2007. 448(7156): p. 894–900.

50. Corbit, V.L., et al., Strengthened inputs from secondary motor cortex to striatum in a mouse model of compulsive behavior. J Neurosci, 2019.

51. Manning, E.E., et al., Impaired instrumental reversal learning is associated with increased medial prefrontal cortex activity in Sapap3 knockout mouse model of compulsive behavior. Neuropsychopharmacology, 2019. 44(8): p. 1494–1504.

52. Taylor, S., Molecular genetics of obsessive-compulsive disorder: a comprehensive meta-analysis of genetic association studies. Mol Psychiatry, 2013. 18(7): p. 799–805.

53. Belanger, M., I. Allaman, and P.J. Magistretti, Brain energy metabolism: focus on astrocyte-neuron metabolic cooperation. Cell Metab, 2011. 14(6): p. 724–38.

54. Jaffe, A.E., et al., Genetic neuropathology of obsessive psychiatric syndromes. Transl Psychiatry, 2014. 4: p. e432.

55. Fromer, M., et al., Gene expression elucidates functional impact of polygenic risk for schizophrenia. Nat Neurosci, 2016. 19(11): p. 1442–1453.

56. Benkelfat, C., et al., Local cerebral glucose metabolic rates in obsessive-compulsive disorder. Patients treated with clomipramine. Arch Gen Psychiatry, 1990. 47(9): p. 840–8.

57. Saxena, S., et al., Localized orbitofrontal and subcortical metabolic changes and predictors of response to paroxetine treatment in obsessive-compulsive disorder. Neuropsychopharmacology, 1999. 21(6): p. 683–93.

58. Remijnse, P.L., et al., Reduced orbitofrontal-striatal activity on a reversal learning task in obsessive-compulsive disorder. Arch Gen Psychiatry, 2006. 63(11): p. 1225–36.

59. Remijnse, P.L., et al., Cognitive inflexibility in obsessive-compulsive disorder and major depression is associated with distinct neural correlates. PLoS One, 2013. 8(4): p. e59600.

60. Reynolds, L.M., et al., Transcriptomic profiles of aging in purified human immune cells. BMC Genomics, 2015. 16: p. 333.

61. Lewis, D.A., The human brain revisited: opportunities and challenges in postmortem studies of psychiatric disorders. Neuropsychopharmacology, 2002. 26(2): p. 143–54.

62. Gerfen, C.R., Basal ganglia, in The Rat Nervous System, G. Paxinos, Editor. 2004, Elsevier: Amsterdam. p. 455–508.

63. Yu, X., et al., Reducing Astrocyte Calcium Signaling In Vivo Alters Striatal Microcircuits and Causes Repetitive Behavior. Neuron, 2018. 99(6): p. 1170–1187 e9.

64. Baxter, L.R., Jr., et al., Caudate glucose metabolic rate changes with both drug and behavior therapy for obsessive-compulsive disorder. Arch Gen Psychiatry, 1992. 49(9): p. 681–9.

65. Saxena, S., et al., Cerebral metabolism in major depression and obsessive-compulsive disorder occurring separately and concurrently. Biol Psychiatry, 2001. 50(3): p. 159–70.

66. Swedo, S.E., et al., Cerebral glucose metabolism in childhood-onset obsessive-compulsive disorder. Revisualization during pharmacotherapy. Arch Gen Psychiatry, 1992. 49(9): p. 690–4.

