## Supplementary figures and images for "Transcriptome alterations are enriched for synapse-associated genes in the striatum of subjects with obsessive-compulsive disorder"

### Supplemental Figure 1

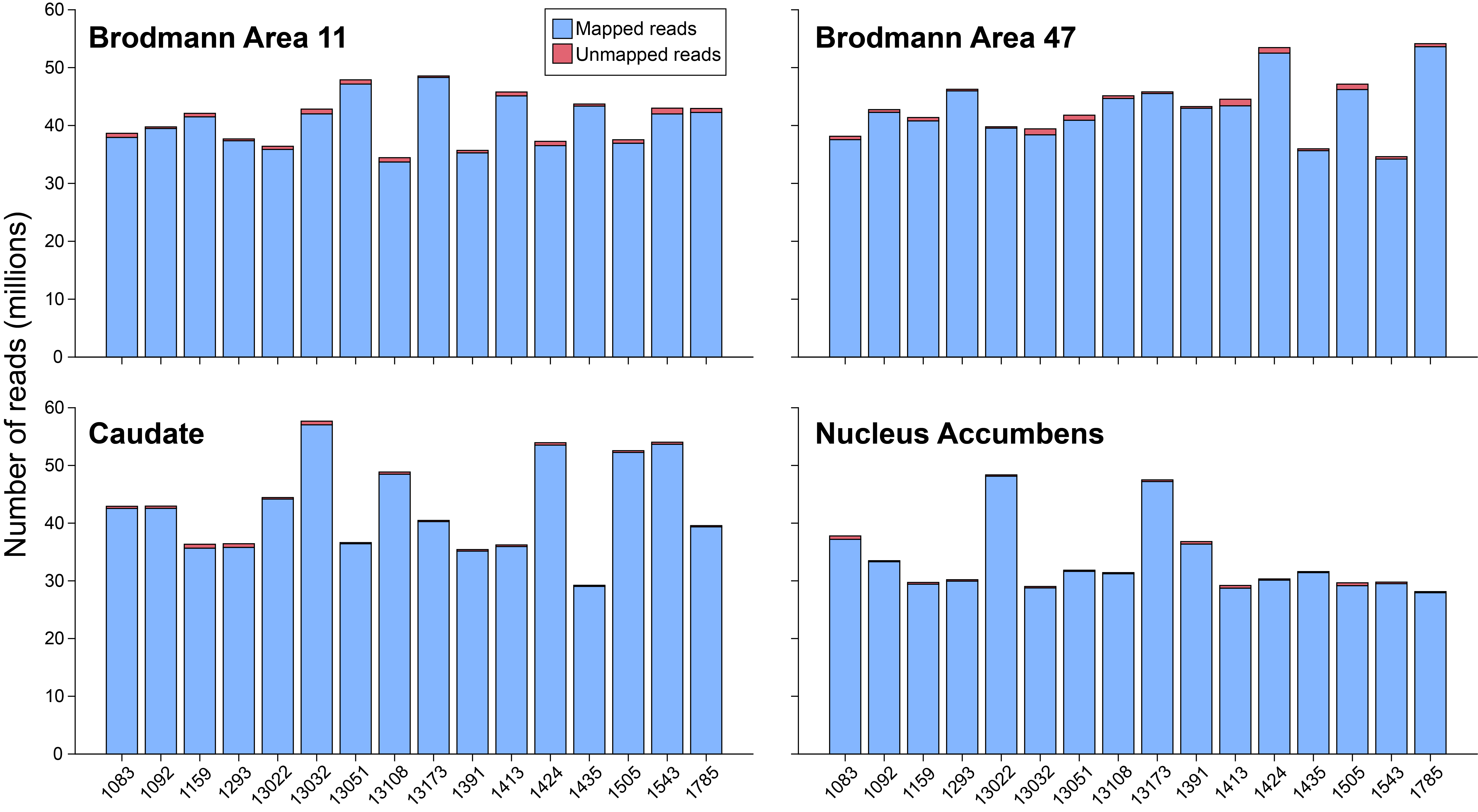

### Supplemental Figure 2

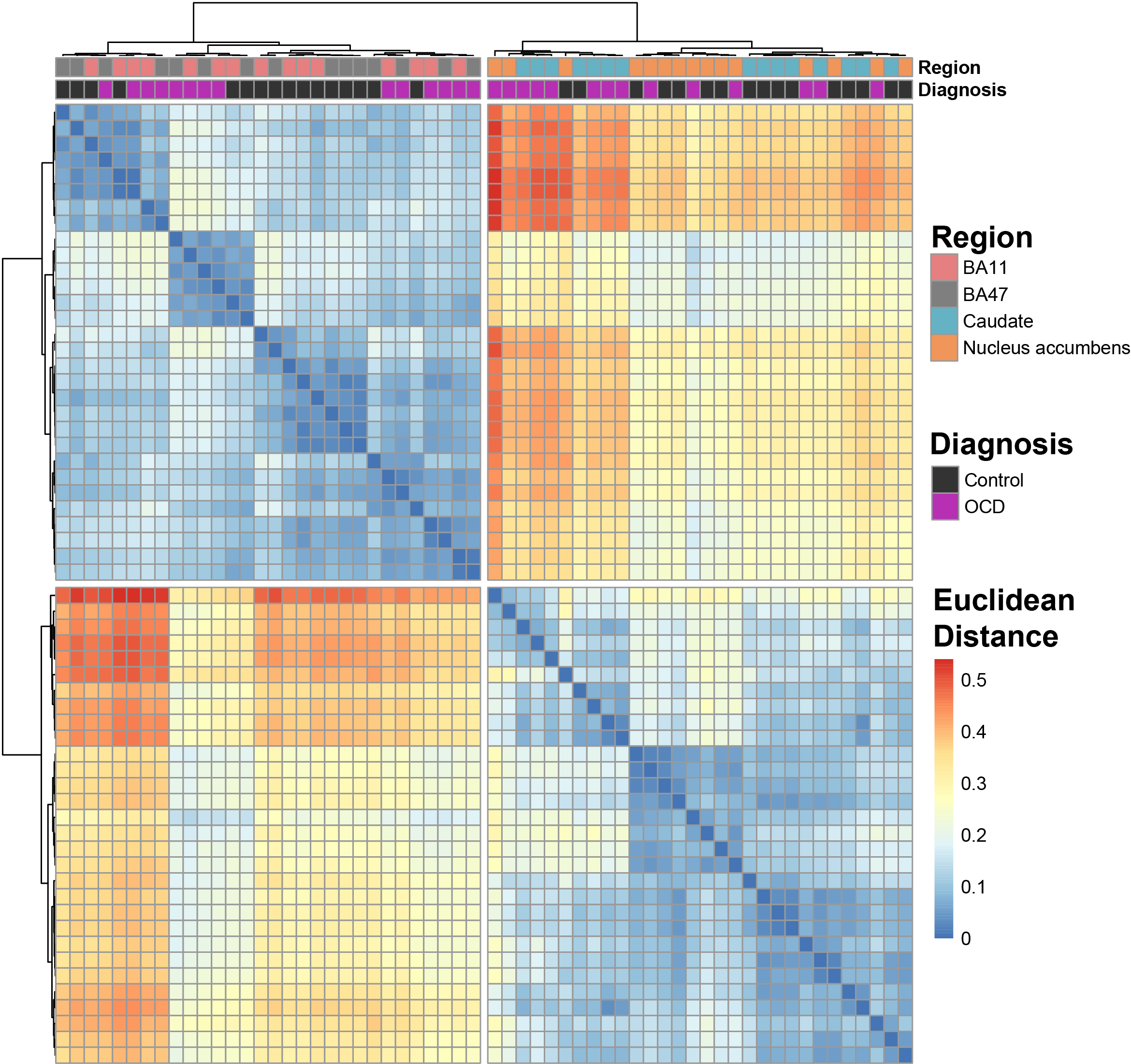

### Supplemental Figure 3

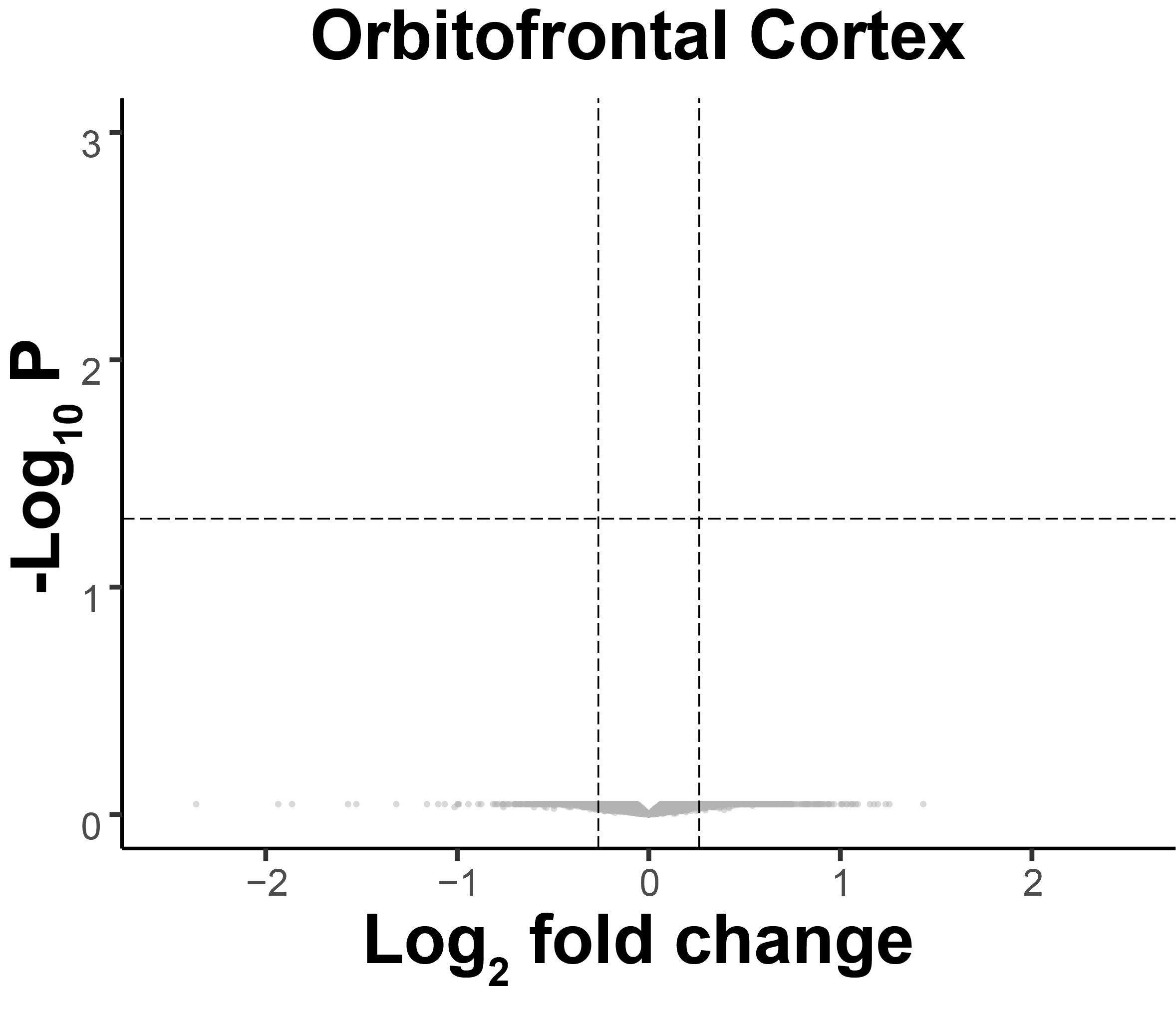
