## Supplement 1 for "Transcriptome alterations are enriched for synapse-associated genes in the striatum of subjects with obsessive-compulsive disorder"

**SUPPLEMENTAL MATERIAL**

**METHODS**

*Tissue Collection & RNA Extraction:* Standardized amounts (50 mm3) of gray matter were collected via dissection of sections cut on cryostat from four separate brain regions identified cytoarchitectonically: medial orbitofrontal cortex (mOFC, BA11), lateral orbitofrontal cortex (lOFC, BA47), head of the caudate nucleus, and nucleus accumbens core. RNA was extracted using an RNeasy Plus Mini kit (QIAGEN, Valencia, CA) as previously described [1]. RNA quality was assessed using RNA ScreenTape Analysis (Agilent Technologies, Santa Clara, CA) and RNA concentration was estimated using a Qubit Fluorometer (ThermoFisher Scientific, Waltham, MA).

*RNA sequencing:* Messenger RNA (mRNA) sequencing was performed using the Illumina NextSeq 500 platform (Illumina Inc, San Diego, CA). Complementary DNA (cDNA) libraries were generated using TruSeq Stranded mRNA Library Preparation kits (Illumina Inc, San Diego, CA) in which poly-adenylated mRNA molecules were purified, fragmented, and then reverse transcribed into first and second cDNA strands. Single-end mRNA sequencing was then performed at a targeted depth of 40 million reads per sample. Reads were aligned to human genome reference sequence GRCh38.p10 and annotated via Ensemble Archive Release v91; the number of reads that mapped (or did not map) to the reference sequence is depicted in Supplemental Figure 1. Quality control (QC) was conducted using CLC Genomics (QIAGEN) and raw read counts were exported for downstream analyses.

*Differential Expression:*	To identify drivers of variation in gene expression, the relationship between gene expression values and covariates – diagnosis, sex, age, post-mortem interval, pH, RNA integrity number, or brain region – were assessed using generalized linear regression [2]. Briefly, following Fromer, Roussos [3], we sought a parsimonious model consisting of covariates that predicted each gene’s expression. To accomplish this goal, we implemented a stepwise forward procedure, beginning with diagnosis and sex as the first predictors in the model. Next, the covariate that maximized the number of genes showing improved fit between the covariate and gene expression, as judged by Bayesian information criterion (BIC), was added to this model. This process of sequentially adding covariates to the model was repeated until only a minority of the gene expression values (<5%) were improved by the next covariate (Supplemental Table 1). Although the relatively small sample size limits efficacy of selecting covariates for predicting variation in gene expression, we believe this process, which includes only covariates that statistically affect transcript expression, is sufficient to obtain a robust model.

To identify genes differentially expressed between OCD subjects and unaffected comparison subjects, either across brain regions or within a single region, we used the covariate model described above. Expression counts were transformed into log2CPM and fitted to covariates using limma. For each gene, the standard deviation of expression was estimated using empirical Bayes techniques to obtain voom precision weights, which were then used to estimate differential expression in this weighted linear regression context [4-6]. Significance of each contrast was determined using Bayes factors, assuming that 1% of genes were expected to be differentially expressed. When analyzing the regions simultaneously, we used the average correlation among regions in gene expression, estimated by limma, to account for repeated observations within subject. Region-specific effects were then estimated simultaneously for all regions, after which significance was judged for appropriate contrasts (FDR-adjusted p-value was set at α=0.05).

*Data Quality Control:* On average, 98.5% of reads were mapped for each sample and the number of mapped reads did not differ significantly between OCD subjects and unaffected comparison subjects (t=1.6; df=14; p=0.14; Supplemental Figure 1). Sequence reads for the four brain regions of the 16 subjects were mapped to 58,243 transcripts, which were further reduced to 18,993 expressed RefSeq genes (v.2015-01). Of those, genes were retained that had at least 1 count per million (CPM) in at least half of the samples for at least one of the four brain regions; and if, over all subjects and brain regions, gene j’s coefficient of variation CVj<T, where T is defined as the mean of CVj over all j plus 3 standard deviations of CV. This resulted in 14,211 genes whose expression passed QC.

To evaluate the extent to which gene expression was distinct for the four brain regions, pairwise consensus correlations were obtained from the limma package in R (v.3.3.19; [2, 6]) and visualized using a heatplot (pheatmap package in R) over all subjects. Pairwise consensus correlations over genes were high between BA11 and BA47 (0.71) compared to other pairwise correlations (0.16-0.25; Supplemental Figure 2). Thus, data for BA11 and BA47 were treated as from a single brain region termed "OFC" by averaging expression, per gene and subject, over BA11 and BA47. In this process, however, we noted one male OCD subject for whom the BA47 and nucleus accumbens samples had likely been inadvertently switched. Thus, this subject was removed from further analysis. After removal, seven OCD subjects and eight unaffected comparison subjects remained. After re-running the QC steps based on the new data, specifically over three brain regions and with one subject removed, there were 14,184 genes remaining for analyses for all brain regions and 13,623, 13,889, and 13,756 for OFC, caudate, and NAcc, respectively.

*Gene Set Enrichment:* To investigate whether differentially expressed genes between OCD subjects and unaffected comparison subjects were enriched for biologically relevant gene-set pathways, we used the Gene Set Enrichment Analysis (GSEA) platform [7]. This platform contains various molecular signature databases (MSigDB, v7.0; [8]), including curated gene sets that are publicly available and biologically relevant. Enrichment was evaluated on the differentially expressed genes from the analysis of all brain regions together and per individual brain region. Within the MSigDB, we focused on the following general gene sets (excluding sets that were curated for specific purposes): Hallmark [50 sets]; the C2 curation (sub-databases included canonical [2,199 sets], KEGG [186 sets], and REACTOME [1,499 sets]); and the C5 Gene Ontology (GO) curation (sub-databases included biological process [7,350 sets], cellular component [1,001 sets], and molecular function [1,645 sets]). For each gene set containing at least 30 genes, differentially expressed genes were analyzed for enrichment using Fisher's exact test and “significance” judged by FDR (for sets with enrichment odds ratio exceeding one).

We determined co-occurrence for the GO gene sets that were significantly enriched in OCD in the global analysis. Genes were defined as co-occurring if present in more than one gene set. A distance value was computed per gene set pair, defined as the fraction of overlapping genes divided by the number of unique genes [9]. A network was computed using the force-directed graph drawing algorithm directed by Fruchterman-Reingold (Figure 2) [10]. Hierarchical clustering was performed on the computed distance values using the Ward D2 method [11]; within the networks, four sub-networks were now distinguishable (Figure 2).

*Cell Type Composition:* The composition of the tissue samples in terms of broad cell type fractions–excitatory neurons, medium spiny neurons (also known as spiny projection neurons), interneurons, astrocytes, oligodendrocytes, ependymal cells, immune cells, and vascular cells – was estimated using deconvolution methods. The principle behind this method is that the bulk expression of a gene for a tissue sample, here measured using RNAseq, is a convolution of the number of cells of each type comprising the sample and the average expression of the gene within the cells of each type. Although the estimates from deconvolution more closely map onto “expression fractions”, we will continue to follow convention and refer to these as “cell type fractions”. To identify a specific feature of this convolution, namely the fraction of each cell type, we utilized a reference transcriptome of single cell-based RNAseq data obtained from mouse cortex and striatum and containing 62,155 cells of various cell types [12]. This reference database was used to derive a “signature matrix” (genes by cell types), which was generated by averaging the observed expression of the top 50 marker genes per cell type over 62,155 cells belonging to the general cell types described above (following [13]). Finally, using the observed gene expression patterns in the region and the signature matrix, non-negative least squares deconvolution estimated the cell type fractions for each brain region and subject in our dataset. Cell type expression profiles of OCD subjects were compared to unaffected comparison subjects using a linear model, adjusting for brain region, sex, age, PMI, pH, and RIN, for the combined brain regions and each region separately.

**LITERATURE CITED**

1.	Piantadosi, S.C., et al., *Lower excitatory synaptic gene expression in orbitofrontal cortex and striatum in an initial study of subjects with obsessive compulsive disorder.* Mol Psychiatry, 2019.

2.	Bates, D., et al., *Fitting Linear Mixed-Effects Models Using lme4.* 2015, 2015. **67**(1): p. 48 %J Journal of Statistical Software.

3.	Fromer, M., et al., *Gene expression elucidates functional impact of polygenic risk for schizophrenia.* Nat Neurosci, 2016. **19**(11): p. 1442-1453.

4.	Law, C.W., et al., *voom: Precision weights unlock linear model analysis tools for RNA-seq read counts.* Genome Biol, 2014. **15**(2): p. R29.

5.	Phipson, B., et al., *Robust Hyperparameter Estimation Protects against Hypervariable Genes and Improves Power to Detect Differential Expression.* Ann Appl Stat, 2016. **10**(2): p. 946-963.

6.	Ritchie, M.E., et al., *limma powers differential expression analyses for RNA-sequencing and microarray studies.* Nucleic Acids Res, 2015. **43**(7): p. e47.

7.	Subramanian, A., et al., *Gene set enrichment analysis: a knowledge-based approach for interpreting genome-wide expression profiles.* Proc Natl Acad Sci U S A, 2005. **102**(43): p. 15545-50.

8.	Liberzon, A., et al., *Molecular signatures database (MSigDB) 3.0.* Bioinformatics, 2011. **27**(12): p. 1739-40.

9.	Ruckdeschel, P., et al., *S4 Classes for Distributions.* R News, 2006. **6(2)**: p. 2-6.

10.	Fruchterman, T.M.J. and E.M. Reingold, *Graph drawing by force-directed placement.* 1991. **21**(11): p. 1129-1164.

11.	Müllner, D., *fastcluster: Fast Hierarchical, Agglomerative Clustering Routines for R and Python.* 2013, 2013. **53**(9): p. 18 %J Journal of Statistical Software.

12.	Zeisel, A., et al., *Molecular Architecture of the Mouse Nervous System.* Cell, 2018. **174**(4): p. 999-1014 e22.

13.	Wang, J., B. Devlin, and K. Roeder, *Using multiple measurements of tissue to estimate subject- and cell-type-specific gene expression.* Bioinformatics, 2020. **36**(3): p. 782-788.
